# Hidden Medicines in Herbarium Collections: Medicinal Plant Diversity and Ethnopharmacological Knowledge from GSUH, Nigeria

**DOI:** 10.64898/2026.10.06.756981

**Authors:** Daniel A. Zhigila, Suleiman D. Abdul, Ibrahim Inuwa, Usman A. Yuguda, Fatima BJ Sawa

**Author notes:** Corresponding author: Daniel A. Zhigila.

## Abstract

Herbarium collections represent an important but frequently underutilized source of information for documenting medicinal plant diversity and preserving ethnomedicinal knowledge. We assessed the medicinal plant diversity represented in herbarium records from the Gombe State University Herbarium (GSUH), Nigeria, and evaluated patterns in taxonomic composition, therapeutic applications, and plant parts reported for medicinal use. We also tested the hypothesis that the plant parts used medicinally are not randomly distributed across therapeutic or ailment categories, but that particular plant parts are disproportionately associated with specific therapeutic applications. The resulting dataset comprised 1,576 specimen records, including 768 recorded as medicinal plant species across 94 families and 1,426 medicinal-use records. Fabaceae was the most species-rich family (15.8%), followed by Asteraceae (9.3%), Malvaceae (4.8%), Lamiaceae (4.4%), Rubiaceae (3.2%), and Apocynaceae (3.2%). The most frequently represented therapeutic themes were infections and sexually transmitted diseases (50.7%), gastrointestinal disorders (41.9%), pain, inflammation, and rheumatism (41.9%), fever and malaria (40.2%), and skin diseases and wounds (40.0%). Individual species had a mean of 2.98 therapeutic themes, with medicinal breadth ranging from 1 to 7 themes. A species-aware permutation test confirmed that the observed plant-part × therapeutic-theme association was significantly stronger than expected under random association (9,999 permutations, *P* < 0.0001). These findings demonstrate the value of herbarium collections as taxonomically anchored resources for recovering information on medicinal plant diversity. Integrating herbarium records with published ethnomedicinal knowledge provides a reproducible framework for linking plant diversity, taxonomy and traditional medicinal applications and highlights the potential of underutilised African herbaria for ethnopharmacological research.

## 1. INTRODUCTION

Medicinal plants remain an important component of healthcare systems worldwide, particularly in regions where traditional medicine provides culturally embedded and locally accessible approaches to disease management (Dalamagka, 2024; Aliaj et al., 2026; Latif and Nawaz, 2026). The World Health Organization recognizes traditional medicine as a major global health resource (WHO, 2025). It emphasizes the need for evidence-based research, appropriate documentation, safety, sustainability, and respect for Indigenous and local knowledge. In sub-Saharan Africa, plant-based remedies are deeply integrated into traditional healthcare, with medicinal species used for a broad spectrum of infectious, inflammatory, gastrointestinal, dermatological, reproductive and chronic conditions (Adetunji, 2026). The region’s medicinal flora is correspondingly extensive: Van Wyk (2020) documented at least 4,576 vascular plant species across 1,518 genera and 192 families as components of Traditional African Medicine.

This diversity represents not only a reservoir of culturally transmitted therapeutic knowledge but also a potentially important source of pharmacologically active natural products. Indeed, the correspondence between ethnobotanical use and biological activity documented for Nigerian medicinal plants illustrates the potential of traditional knowledge to inform natural-product research and drug discovery (Lifongo et al., 2014).

The relationship between medicinal-plant diversity and traditional knowledge, however, is not simply a consequence of plant availability. Patterns of medicinal-plant selection can reflect ecological availability, cultural preferences, therapeutic traditions and, in some cases, the occurrence of particular classes of secondary metabolites (Adetunji, 2026). Several studies (e.g., van Wyk, 2020; Plaatjie et al., 2024; Adetunji, 2026) revealed that several plant families are disproportionately represented in traditional African medicine, with Fabaceae, Rubiaceae, Asteraceae, Malvaceae and Euphorbiaceae among the most species-rich groups. At the same time, medicinal-plant knowledge is dynamic and geographically heterogeneous, with different cultural groups developing distinctive materia medica and therapeutic practices (Saslis-Lagoudakis et al., 2014). Recent inventories demonstrate that substantial numbers of medicinal species continue to be added as previously undocumented knowledge is recovered; for example, Cornelius and Van Wyk (2024) identified 3,784 medicinal species in southern Africa and reported more than 1,200 species not included in the earlier regional inventory. These findings highlight an important methodological issue in ethnopharmacology: apparent differences in medicinal-plant diversity may partly reflect differences in the extent, accessibility and taxonomic quality of documentation rather than differences in actual knowledge or plant use.

Nigeria provides an especially important context for addressing this documentation problem because of the country’s exceptional floristic and cultural diversity (Zhigila et al., 2026). Medicinal plants are used across its major ecological and cultural regions, but the available knowledge remains fragmented among ethnobotanical surveys, local publications, theses, traditional knowledge systems, and biological collections. A recent taxonomic assessment of Nigerian ethnobotanical research compiled 963 plant species belonging to 144 families from 79 studies conducted between 1964 and 2024, while also identifying substantial regional differences in the representation of plant-use knowledge (Rafiu et al., 2025). Such fragmentation is particularly consequential in northern Nigeria, where medicinal knowledge is closely associated with local plant resources and where historical documentation has often been geographically and taxonomically uneven. Importantly, herbarium specimens provide an independent and verifiable foundation for ethnopharmacological research by linking medicinal-use information to taxonomically authenticated plant material, collection localities, and permanently archived voucher specimens, thereby allowing plant identity and the geographical context of reported uses to be independently reassessed (Culley, 2013; Brinckmann et al., 2017; Souza and Hawkins, 2017; Hart et al., 2026). Earlier Nigerian ethnobotanical research already demonstrated the value of voucher-based documentation, in which botanical names, collection numbers, localities and reported medicinal applications were explicitly associated with herbarium specimens (Nwosu, 1998).

Herbarium collections are therefore more than repositories of dried specimens; they are durable biological archives that preserve plant identities, taxonomic relationships, geographical occurrence, and historical patterns of use for subsequent verification and re-evaluation (Davis et al., 2024). In ethnopharmacology, such records provide a taxonomically reliable basis for linking medicinal uses to particular species and can reveal therapeutic applications, plant parts harvested, and taxonomic groups disproportionately represented in medicinal plant repertoires (Souza & Hawkins, 2017; Van Wyk, 2020). This information has both pharmacological and conservation relevance because harvesting roots, bark, bulbs, rhizomes, or whole plants can be more destructive than harvesting renewable organs such as leaves, flowers, fruits, or seeds, particularly in regions where medicinal plants are already threatened by habitat loss and unsustainable harvesting (Schippmann et al., 2002; Moyo et al., 2015; Williams et al., 2019).

Against this background, the Gombe State University Herbarium (GSUH) represents an important but previously underutilised source of medicinal-plant and ethnomedicinal information from northeastern Nigeria. This study uses herbarium-associated records to characterise the medicinal plant diversity documented in the region and to assess the therapeutic dimensions of this knowledge. Specifically, we quantify the taxonomic composition of medicinal plants, assess the representation of plant families, characterise major therapeutic and ailment categories, evaluate the breadth of therapeutic uses associated with individual species, and document the plant parts used in traditional medicine. We also hypothesized that plant parts are not randomly associated with therapeutic applications, but that particular plant organs are disproportionately represented in specific therapeutic categories. Demonstrating such associations would provide a quantitative basis for interpreting patterns of medicinal plant use and identifying plant-use combinations that may be of particular ethnobotanical or conservation interest. By treating herbarium records as structured ethnopharmacological evidence rather than simply as species lists, the study seeks to provide a reproducible baseline for understanding the medicinal flora of Africa using the records at the GSUH and to identify taxonomic and therapeutic patterns that can inform subsequent pharmacological, phytochemical and conservation research. More broadly, the study demonstrates how the mobilisation and analysis of regional herbarium collections can recover geographically embedded medicinal-plant knowledge and make it available for contemporary ethnopharmacological research, while retaining the botanical evidence required for future verification and reinterpretation.

## 2. METHODOLOGY

### 2.1. Herbarium specimens

Herbarium specimens deposited at the Gombe State University Herbarium (GSUH) were examined as the primary source of data for documenting medicinal plant diversity. Each specimen was assessed for taxonomic identity, physical condition, diagnostic morphological characters, and the completeness of label information, including scientific name, collector and collection number, collection date, locality, habitat, and associated collection notes. Specimens with incomplete, synonymised, or uncertain identifications were re-examined using diagnostic morphological characters and, where necessary, relevant regional floras, taxonomic revisions, monographs, and other authoritative literature (e.g., Hutchinson and Dalziel 1972; Akobundu and Agyakwa 2008; Zhigila et al., 2023). Specimens that could not be reliably identified to species were assigned to the highest taxonomic rank supported by the available evidence, at least to genus level, and flagged for further verification. Particular attention was given to resolving closely related taxa, outdated names, and apparent typographical or transcription errors.

### 2.2. Taxonomic verification and nomenclatural standardisation

Taxonomic names were verified using published taxonomic treatments, including Keay 1963; Hutchinson and Daziel 1972; Akobundu and Agyakwa 2008; Ogbudemi 2008; Zhigila et al. (2023), and authoritative nomenclatural databases, mainly Plants of the World Online (POWO, 2026), maintained by the Royal Botanic Gardens, Kew, and the International Plant Names Index (IPNI, 2026) and JSTOR (2006). A master list of taxa represented in the collection was compiled and standardised. Typographical errors, spelling inconsistencies, author-citation differences, and formatting inconsistencies were corrected prior to data analyses. Each taxon was assessed according to its nomenclatural status in POWO. Names recognised as accepted were retained as the standard taxonomic names, whereas synonyms were linked to their corresponding accepted names. Unresolved, ambiguous, or unrecognised names were flagged and investigated further using relevant taxonomic literature listed above. The final taxonomic backbone was standardised at family, genus, and species levels. This process minimised duplicate representation of taxa arising from nomenclatural variation and provided a consistent framework for quantifying medicinal plant diversity and linking herbarium records with published ethnomedicinal information.

### 2.3. Metadata capture, cleaning and database management

Information from specimen labels was transcribed into a structured digital database. Medicinal-use information obtained from the literature was maintained separately from specimen metadata to distinguish voucher-derived information from literature-based ethnomedicinal evidence. Scientific names and taxonomic terminology were standardised during data entry to minimise nomenclatural inconsistencies and facilitate data retrieval, analyses, and subsequent dissemination (Hedrick et al., 2020; Borsch et al., 2020; Bräuchler et al., 2021). Locality information was assessed for geographic consistency and, where coordinates were absent but sufficiently detailed locality descriptions were available, specimens were georeferenced using appropriate geographic reference resources. The resulting dataset was therefore a curated, specimen-linked dataset suitable for subsequent taxonomic and ethnomedicinal analyses.

### 2.5. Medicinal plant assessment and ethnomedicinal information

The medicinal status and reported uses of the identified taxa were assessed through a targeted review of published ethnobotanical, ethnomedicinal, pharmacognostic, and medicinal-plant literature, including Odugbemi (2008), Zhigila et al. (2023), Adeniran and Akindele (2024), Rafiu et al. (2025), and other relevant literature. For each validated taxon, published evidence was examined to determine whether medicinal uses had been documented and to extract information on the reported therapeutic application and plant part used. Medicinal uses were based exclusively on published evidence and personal communication with local herbal sellers in Gombe and were not inferred from taxonomic identity. Where multiple sources reported uses for the same species, the information was consolidated under the accepted species name while retaining distinct therapeutic applications and plant parts reported across sources.

Reported plant parts were standardised into comparable categories, including leaves, roots, bark, stems or shoots, flowers, fruits, seeds, whole plants, underground organs, and exudates. Individual medicinal applications were similarly grouped into broader therapeutic or ailment themes to facilitate quantitative comparisons among species. The original reported uses were retained in the database, while broader categories were used only for statistical summarisation and visualisation.

### 2.6. Statistical analysis

To test whether plant parts were non-randomly associated with therapeutic applications, medicinal-use records were standardized and organized into species–plant-part–therapeutic-theme associations. Repeated records for the same species, plant part and therapeutic theme were consolidated so that multiple literature records did not disproportionately influence the analysis. Plant parts were grouped into standardized categories, and medicinal uses were assigned to broader therapeutic or ailment themes. A plant-part × therapeutic-theme contingency table was constructed and tested using Pearson’s chi-square test of independence, with Cramér’s V calculated as a measure of association strength. Standardized residuals were examined to identify combinations occurring more or less frequently than expected under independence and were visualized as a residual heatmap. As individual species could contribute to multiple plant-part and therapeutic-theme combinations, a species-aware permutation test with 9,999 permutations was additionally used to assess the robustness of the observed association while accounting for the non-independence of observations. Correspondence analysis was used as an exploratory multivariate approach to visualize the major associations between plant-part categories and therapeutic themes. All statistical analyses and graphical representations were performed in R version 4.6.2 (R Core Team, 2026).

## 3. RESULTS

### 3.1 What are the taxonomic and ethnomedicinal composition?

Of the 768 unique vascular plant species represented in the GSUH collection, 475 (61.8%) had documented medicinal uses, whereas the remaining 293 species (38.2%) were documented for other uses or lacked evidence of medicinal use. The GSUH medicinal plant species consist of 94 plant families, with 1,426 species–ailment records assigned to 16 broad ailment or therapeutic themes. Species representation was strongly uneven among families. Fabaceae was the most species-rich family, comprising 75 medicinal species, followed by Asteraceae (44 species), Malvaceae (23 species), Lamiaceae (21 species), and Rubiaceae and Apocynaceae (15 species each). Together, these six families accounted for 40.6% (n=193) of the medicinal species represented in the dataset. The remaining families contributed with fewer species, indicating a pronounced concentration of medicinal-plant diversity within a relatively small number of taxonomic groups (Fig. 1).

**Figure 1.**
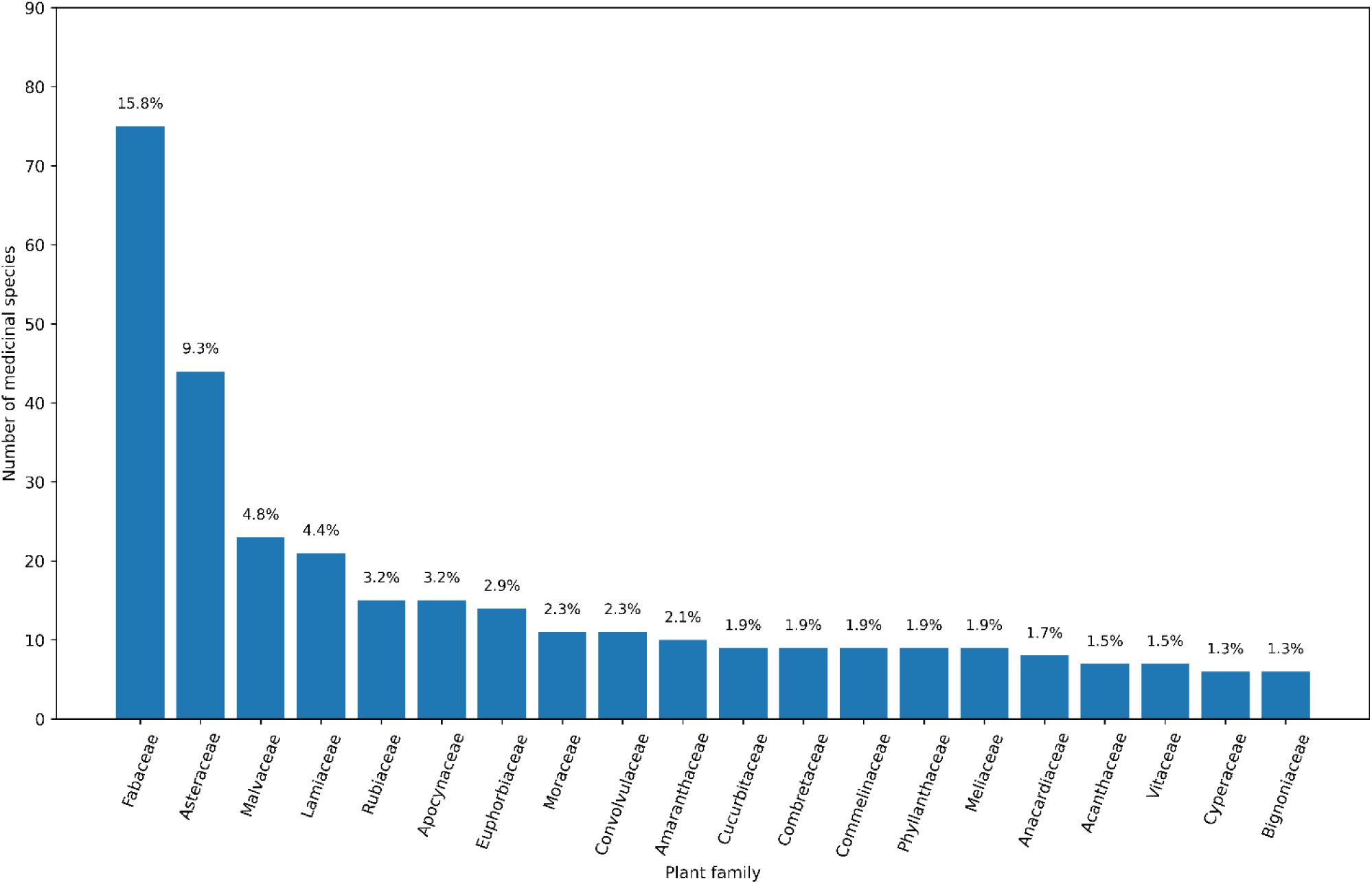
Species richness of the medicinal flora represented in the Gombe State University Herbarium (GSUH), showing the 20 most species-rich plant families. Bars represent the number of unique medicinal species recorded in each family, while percentages indicate the contribution of each family to the total medicinal species dataset (*n* = 475).

The distribution of medicinal species across therapeutic themes was similarly uneven (Fig. 2A). Infections and sexually transmitted diseases represented the largest thematic category, involving 241 species (50.7% of all medicinal species). This was followed by gastrointestinal disorders, with 199 species (41.9%), pain, inflammation and rheumatism, also with 199 species (41.9%), fever and malaria, 191 species (40.2%), and skin diseases and wounds, involving 190 species (40.0%). Respiratory conditions accounted for 101 species (21.3%). Other therapeutic categories were represented by substantially fewer species, including diabetes and metabolic disorders (44 species; 9.3%), cardiovascular and blood-related conditions (38; 8.0%), eye, ear and oral conditions (34; 7.2%), liver disorders and jaundice (32; 6.7%), snake and insect bites or poisoning (30; 6.3%), neurological and sleep-related conditions (29; 6.1%), urinary and kidney disorders (29; 6.1%), reproductive and fertility-related conditions (28; 5.9%), parasites and worms (21; 4.4%), and cancer and tumours (10; 2.1%).

**Figure 2.**
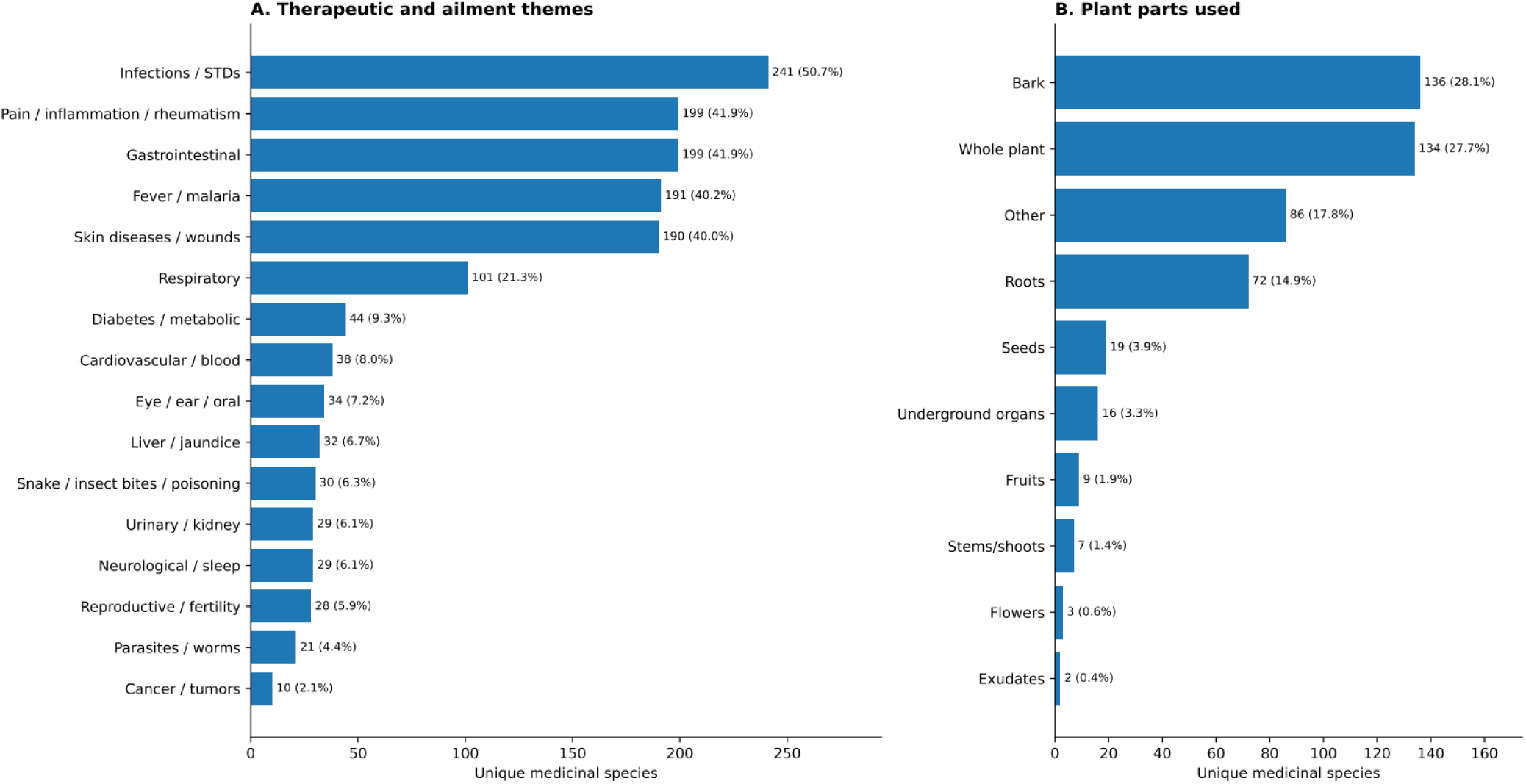
Therapeutic applications and plant parts used in medicinal species. (A) Number of unique medicinal plant species associated with each of 16 therapeutic or ailment themes. Percentages are relative to the 475 unique medicinal species; because individual species may have multiple applications, percentages are not expected to sum to 100%. (B) Number of unique medicinal species associated with broad plant-part categories. Percentages are relative to the 484 species represented in the plant-part dataset, and categories are not mutually exclusive because a species may have more than one medicinally used part.

### 3.3 What plant parts are harvested?

Multiple plant parts were recorded for some species (Fig. 2B). When the reported plant parts were standardized into broad categories, whole plants (28.1) and leaves (27.7%) were among the most frequently represented plant parts, followed by roots (17.8%), bark (14.9%, seeds (3.9%), fruits (1.9%), and stems or shoots (1.4%) flowers (0.6%) and exudates (0.4%) (Fig. 2). The occurrence of multiple plant parts for individual species indicates that medicinal use was not restricted to a single morphological structure in a substantial proportion of the documented taxa.

### 3.2 What is the breadth of medicinal use?

Medicinal species varied considerably in the breadth of ailments for which they were recorded (Fig. 3). Across the species recorded, each species was associated with an average of approximately three therapeutic themes, with a median of three themes per species. The distribution was right-skewed, with many species associated with a relatively small number (1.3%) of therapeutic categories and a smaller number of species recorded for multiple categories (37.7%) (Fig. 3). The most broadly used species were associated with up to seven different ailment or therapeutic themes. Thus, the medicinal flora at GSUH comprised both relatively specialised species and species with broad ethnomedicinal applications.

**Figure 3.**
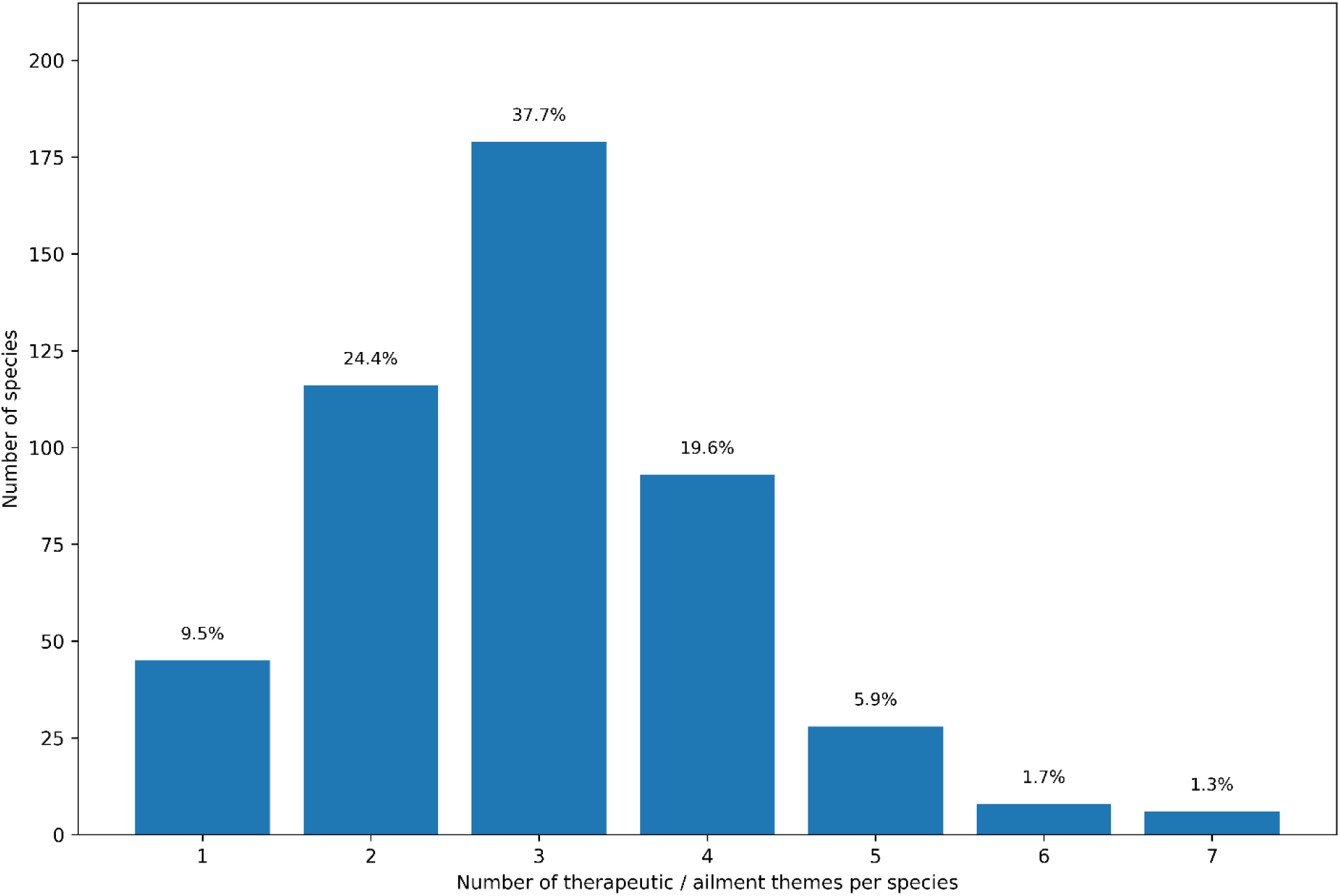
Distribution of medicinal plant species according to the number of distinct therapeutic or ailment themes for which medicinal uses have been documented. Bars represent the number of species, and percentages indicate their proportion of the total medicinal species dataset (*n* = 475).

### 3.4 Does plant-part selection correspond to therapeutic application?

The distribution of plant parts across therapeutic applications was significantly non-random, supporting our hypothesis that particular plant parts are disproportionately associated with specific therapeutic categories (Table 1). Across the 475 medicinal species represented in both the plant-part and therapeutic-use datasets, 2,351 unique species–plant-part–therapeutic-theme associations were identified. The plant-part × therapeutic-theme contingency table showed a significant association (χ² = 294.93, df = 150, *P* < 0.001), although the overall effect size was modest (Cramér’s V = 0.112). Importantly, a species-aware permutation test, which preserved the observed multiplicity of plant parts and therapeutic applications at the species level (Fig. 4), confirmed that the observed association was substantially stronger than expected under random association (9,999 permutations; *P* < 0.0001).

**Table 1:** Statistical summary of medicinal plant diversity and plant-part × therapeutic-theme associations.

| Statistic | Values |
| --- | --- |
| Species represented in both datasets | 475 |
| Unique species–part–theme associations | 2,351 |
| Pearson $\chi^2$ | 294.93 |
| df | 150 |
| Conventional $\chi^2$ P-value | <0.001 |
| Cramér's V | 0.112 |
| Species-aware permutations | 9,999 |
| Species-aware permutation P | <0.0001 |
| CA Dimension 1 | 42.7% inertia |
| CA Dimension 2 | 24.6% inertia |
| First two CA dimensions | 67.3% of inertia |

**Figure 4.**
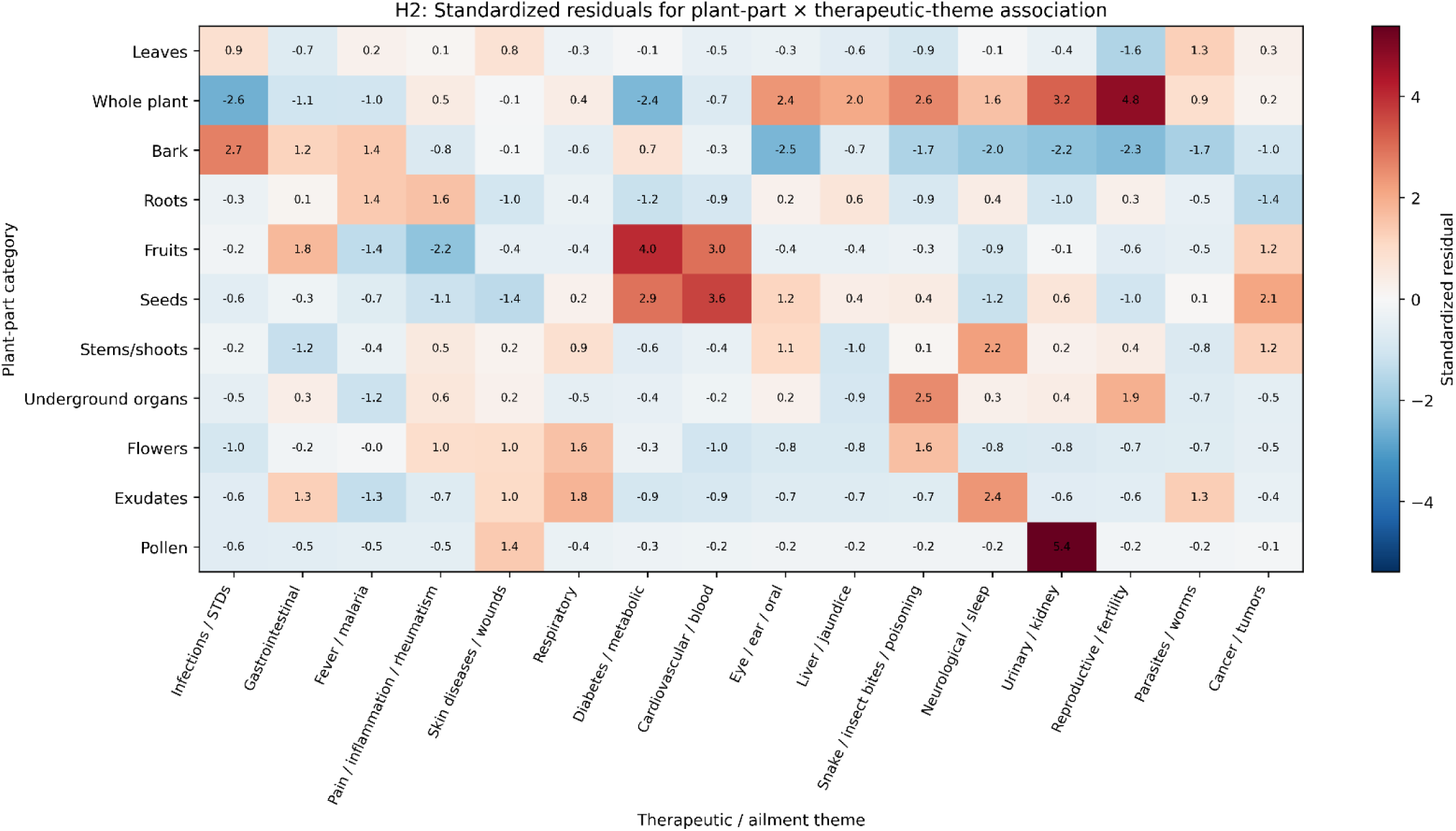
Non-random association between plant parts and therapeutic applications. Heatmap of standardized residuals from the plant-part × therapeutic/ailment-theme contingency analysis. Positive residuals indicate combinations occurring more frequently than expected under independence, whereas negative residuals indicate combinations occurring less frequently than expected.

Standardized residuals revealed several disproportionately frequent combinations, including flowers with urinary/kidney applications, whole plants with reproductive/fertility applications, fruits with diabetes/metabolic applications, seeds with cardiovascular/blood applications, and bark with infections/sexually transmitted diseases. Correspondence analysis further summarized the multivariate structure of these associations, with the first and second dimensions explaining 42.7% and 24.6% of the inertia, respectively, together accounting for 67.3% of the total inertia.

The cross-tabulation of plant families and therapeutic themes showed that the major species-rich families contributed to several therapeutic categories rather than being restricted to individual ailments (Fig. 5). Fabaceae, Asteraceae, Malvaceae, Lamiaceae and other species-rich families were represented across multiple major therapeutic themes. Conversely, several less species-rich families contributed comparatively few species to individual therapeutic categories. This pattern demonstrates that both taxonomic composition and therapeutic composition were heterogeneous across the GSUH medicinal-plant records. When source-record frequency was compared with the number of unique species, the ranking of therapeutic categories was broadly consistent, although the number of records was generally greater than the number of unique species because individual species could be associated with multiple records within the dataset. Consequently, analyses based on unique species provide a more conservative measure of the taxonomic breadth of medicinal knowledge, whereas record counts reflect the frequency and multiplicity of documented species, i.e., ailment associations (Fig. 5).

**Figure 5.**
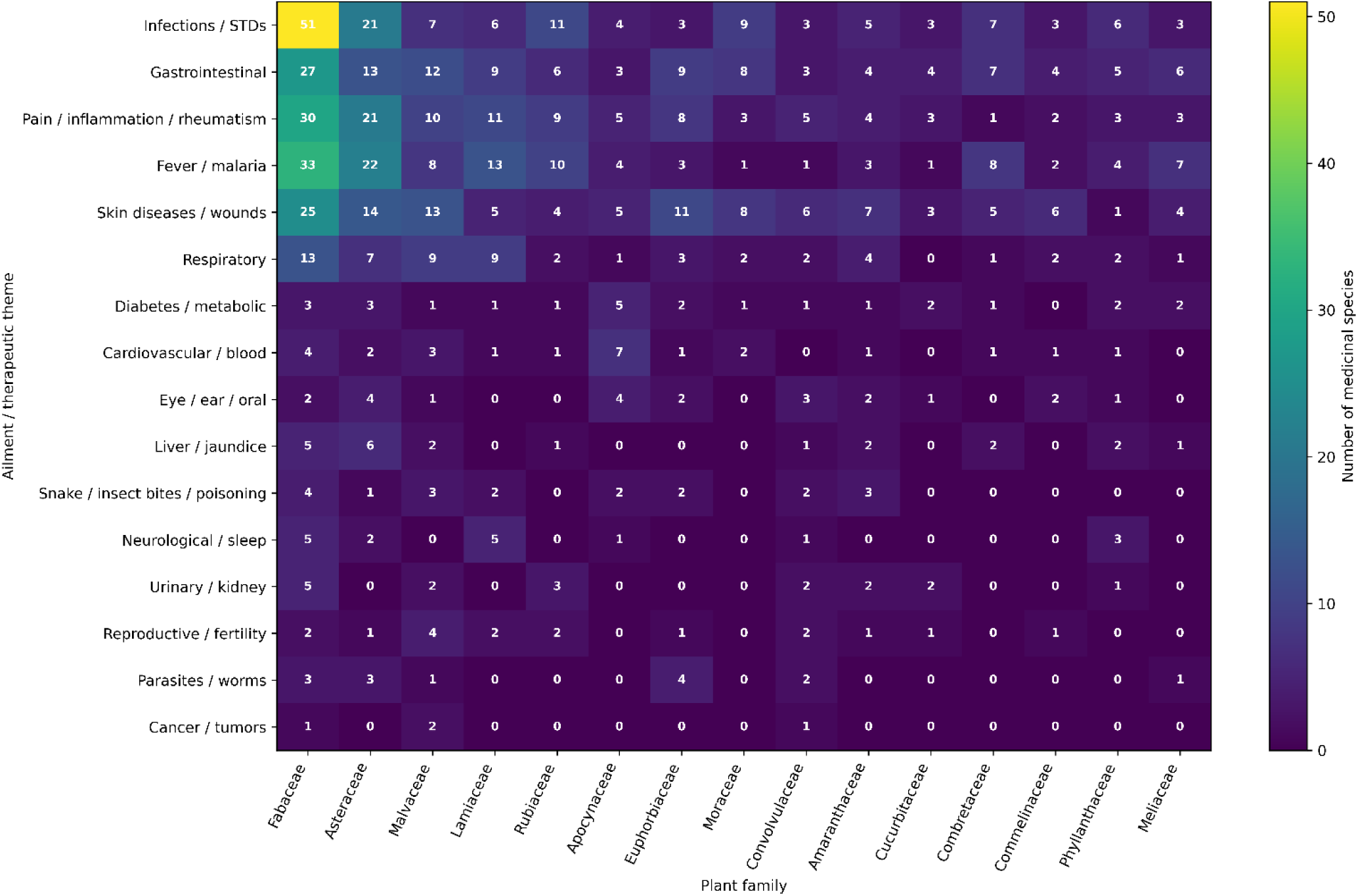
Heatmap indicating the number of unique medicinal plant species associated with each combination of the 15 most species-rich plant families and therapeutic or ailment themes. Cell values represent the number of medicinal species documented for each family.

## 4. Discussion

This study demonstrates that herbarium records can provide a substantial, taxonomically anchored source of ethnomedicinal knowledge for an underdocumented region of Nigeria. The Gombe State University Herbarium (GSUH) records comprised 475 medicinal plant species distributed across 94 families and associated with 16 broad therapeutic or ailment categories. This level of diversity reinforces the importance of plant-based medicine within local healthcare practices and places the medicinal flora documented at GSUH within the broader pattern of exceptionally high medicinal-plant diversity reported for Africa (Adeniji 2026). This supports the findings of Van Wyk (2020), who estimated that at least 4576 vascular plant species are used in Traditional African Medicine, while Moyo et al. (2015) highlighted both the extensive medicinal-plant heritage of sub-Saharan Africa and the increasing pressures threatening its persistence. More recently, a systematic review of Nigerian ethnobotanical literature identified 963 useful plant species belonging to 144 families, with medicinal applications representing the dominant category of plant use (Rafiu et al., 2025). The substantially different totals between the national literature synthesis and the present GSUH inventory should not be interpreted as contradictory estimates of Nigerian medicinal-plant diversity. Rather, they reflect differences in geographic coverage, data sources, sampling intensity, documentation history, and the distinction between literature-derived ethnobotanical records and voucher-associated herbarium records. The present findings therefore demonstrate the value of regional collections for revealing medicinal-plant diversity that may remain obscured in national-scale compilations.

The taxonomic structure of the GSUH medicinal flora provides further insight into the plants that contribute disproportionately to the local medicinal repertoire. The prominence of Fabaceae is particularly consistent with broader African patterns (Sawa et al., 2025). Van Wyk (2020) found that Fabaceae was strongly represented among African medicinal plants and suggested that its prominence cannot be explained solely by the high floristic richness of the family; ecological availability, cultural selection and phytochemical characteristics may all contribute to its medicinal prominence. Fabaceae was also the most represented family in the recent synthesis of Nigerian ethnobotanical research, with 127 species recorded across the reviewed literature (Rafiu et al., 2025). The representation of Asteraceae, Malvaceae, Lamiaceae, Rubiaceae and Apocynaceae in the GSUH records similarly agrees with the recurrent importance of these families in African traditional medicine. Nevertheless, family-level prominence should not be equated with pharmacological superiority. The number of medicinal species within a family reflects, among other factors, the size of the family, local abundance, accessibility, cultural preferences and historical intensity of ethnobotanical documentation. Consequently, the strong representation of these families identifies important taxonomic patterns for further investigation but does not by itself establish therapeutic effectiveness.

The therapeutic profile of the GSUH collection indicates that medicinal plants are employed to address a remarkably broad spectrum of health concerns. Infections and sexually transmitted diseases represented the largest therapeutic category, involving 241 species (50.7% of the medicinal flora), followed by gastrointestinal disorders and pain, inflammation and rheumatism (199 species each), fever and malaria (191 species), and skin diseases and wounds (190 species). These patterns are broadly compatible with the health priorities repeatedly identified in Nigerian ethnobotanical research, where malaria, gastrointestinal disorders, skin conditions, diabetes and other infectious or chronic diseases feature prominently (Zhigila et al., 2023; Rafiu et al., 2025). The high representation of plants used for infections, fever and malaria is particularly noteworthy in an ethnopharmacological context because infectious diseases have historically provided an important pathway through which traditional plant-use knowledge has informed pharmacological investigation. However, the present data describe reported traditional uses rather than experimentally demonstrated therapeutic activity. Consequently, the frequency with which a species is associated with a particular ailment should be interpreted as an indicator of ethnomedicinal importance or knowledge prominence, not as evidence of clinical efficacy, safety or pharmacological potency. The concentration of records in several major therapeutic categories nevertheless provides a useful basis for prioritising species and taxa for subsequent phytochemical, pharmacological and toxicological investigation.

Further, an important finding is the breadth of therapeutic knowledge associated with individual species. On average, each medicinal species was associated with approximately three therapeutic themes (mean = 2.98; median = 3), with individual species spanning between one and seven categories. This pattern indicates that medicinal plants in the GSUH records are not generally restricted to single therapeutic applications; instead, many taxa are embedded within multifunctional systems of traditional healthcare. Such therapeutic versatility may arise from multiple factors, including perceived efficacy across different conditions, availability and accessibility of particular species, cultural transmission of established remedies, and the presence of chemically diverse constituents (Odugbemi, 2008). Van Wyk (2020) argued that the selection of African medicinal plants reflects both ecological availability and non-random cultural and chemical criteria. The repeated use of particular taxa across unrelated therapeutic categories may therefore identify species of particular interest for ethnopharmacological research. At the same time, therapeutic breadth should be interpreted cautiously because a species associated with several ailments may reflect cultural attribution rather than multiple pharmacological activities. Further investigation combining ethnobotanical use data with phytochemistry, bioassays and toxicological assessment would be necessary to distinguish genuine biological versatility from patterns generated by cultural or historical transmission of knowledge.

The pattern of plant parts used has important implications for both pharmacological interpretation and conservation. Leaves and whole plants were among the most frequently represented plant-part categories in the GSUH records, together with roots, bark, seeds, fruits, flowers, and stems or shoots. The use of different plant organs is ethnopharmacologically significant because chemical composition can vary substantially among organs and developmental stages, while harvesting practices have markedly different consequences for plant survival. Harvesting renewable aerial parts such as leaves, fruits and, where reproductive output is not substantially affected, flowers is generally less destructive than harvesting roots, underground organs or bark, which can severely impair plant function, reduce regeneration, or cause mortality, particularly when harvesting is intensive or repeated (Chen et al., 2016; Chamberlain et al., 2019). The use of roots and bark in traditional medicine has consequently been recognised as a potential conservation concern for medicinal plants in sub-Saharan Africa (Moyo et al., 2015). The GSUH plant-part records therefore have value beyond documenting traditional practice: they provide a basis for identifying taxa for which medicinal use may intersect with harvesting pressure. Nevertheless, plant-part frequency in the present dataset should not be interpreted directly as a measure of harvesting intensity, because the number of documented uses does not necessarily correspond to the frequency or quantity of material actually harvested in local communities. Quantitative assessments of harvesting pressure would require field-based studies incorporating use frequency, quantities harvested, harvesting season, plant abundance and regeneration.

The distinction between individual species and source records is also important when interpreting the magnitude of medicinal-plant use. The dataset contained 1,426 species–ailment records but only 475 unique medicinal species, demonstrating that individual species can occur repeatedly because they are associated with multiple therapeutic categories. Failure to distinguish these two levels could substantially exaggerate estimates of medicinal-plant diversity or distort the relative importance of particular therapeutic categories. The present approach, which analyses both unique species and their associated therapeutic records, provides a more informative representation of ethnomedicinal structure. It also allows the identification of species that recur across several therapeutic categories and families that contribute disproportionately to the medicinal flora. This is particularly relevant in Nigeria, where ethnobotanical information remains fragmented across geographically and culturally distinct studies (Rafiu et al., 2025). A standardised, voucher-based approach offers an opportunity to integrate otherwise disconnected observations while retaining the botanical evidence necessary to verify species identities.

The herbarium basis of this study is particularly important in this regard. Voucher specimens provide a permanent physical reference against which plant identifications can be checked, updated and linked to subsequent taxonomic, phytochemical or pharmacological research (Meineke et al., 2018). This is a major advantage over ethnomedicinal reports in which vernacular names or unvouchered species identifications may make subsequent verification difficult. The recent Nigerian ethnobotanical synthesis itself emphasised the importance of taxonomic verification and used resources including Plants of the World Online, GBIF and World Flora Online to standardise plant names (Rafiu et al., 2025). GSUH records extend this principle by retaining the underlying botanical specimens. Such collections are particularly valuable because plant names, family circumscriptions and species concepts change through time; a physical voucher therefore allows historical ethnomedicinal records to be reinterpreted in the light of future taxonomic revisions. Herbarium collections consequently have an important role not only in biodiversity documentation but also in establishing reproducible links between traditional knowledge and contemporary pharmacological research.

The findings also have implications for conservation and the preservation of ethnomedicinal knowledge. Medicinal plants constitute a biologically and culturally valuable component of African plant diversity, yet both the plants and the knowledge associated with them may be vulnerable to environmental change, habitat transformation, overharvesting and generational loss (Moyo et al., 2015). The concentration of medicinal uses within a relatively small number of taxonomic families, combined with the occurrence of highly versatile species, suggests that changes affecting particular plant groups could have disproportionate consequences for local medicinal resources. Conversely, documenting medicinal species in herbaria creates a durable archive that can support future assessments of distribution, conservation status and population change. The GSUH collection can therefore be viewed as both a biodiversity repository and a knowledge infrastructure linking plant taxonomy, traditional healthcare and conservation. This perspective is particularly relevant to Nigeria, where recent synthesis has identified insufficient documentation of indigenous plant-use knowledge as a continuing challenge (Rafiu et al., 2025).

Several limitations should nevertheless be considered. First, the present analysis is based on herbarium-associated records and therefore reflects documented rather than exhaustive local medicinal knowledge. Species absent from the collection cannot be assumed to be absent from local medicinal practice. Second, the therapeutic categories used here simplify complex traditional concepts of disease, diagnosis, and treatment into broader analytical categories, and some individual uses may reasonably fit more than one category. Third, frequency of documentation does not provide a direct measure of use frequency, cultural importance or therapeutic effectiveness. Ethnomedicinal reports should also not be interpreted as evidence of safety or efficacy without appropriate pharmacological and toxicological validation. These limitations are not weaknesses unique to GSUH but represent broader challenges in converting culturally and contextually embedded medicinal knowledge into standardised datasets suitable for comparative analysis. Future work should therefore combine the herbarium inventory with community-based ethnobotanical studies, quantitative indices of cultural importance, phytochemical profiling, pharmacological assays, toxicity evaluation and conservation assessments.

## 5. Conclusion

The GSUH records reveal a rich and taxonomically diverse medicinal flora in Nigeria and demonstrate the value of herbarium collections as repositories of ethnopharmacological knowledge. The occurrence of 475 medicinal species across 94 families, the strong representation of Fabaceae and several other recurrent medicinal families, the concentration of uses in infectious, gastrointestinal, inflammatory, febrile and dermatological conditions, and the substantial therapeutic breadth of individual species collectively indicate a complex medicinal-plant system rather than a small set of isolated remedies. Importantly, the study provides a foundation for moving from documentation towards hypothesis-driven ethnopharmacological research. Species repeatedly associated with multiple therapeutic categories, taxa representing culturally important medicinal families, and plants involving potentially high-impact harvesting practices can be prioritised for targeted pharmacological, phytochemical and conservation studies. Our findings supported the hypothesis that medicinal plant parts are non-randomly associated with therapeutic applications. The significant association demonstrates that particular plant organs are disproportionately represented in specific therapeutic categories, highlighting structured patterns within the ethnomedicinal knowledge documented in the GSUH collection. More broadly, the findings demonstrate that regional herbaria such as GSUH can play a critical role in recovering, validating and preserving medicinal-plant knowledge while simultaneously strengthening the taxonomic foundation upon which contemporary ethnopharmacological research depends.

## Data availability

The underlying specimen and medicinal-use data generated and curated during this study will be made available through an appropriate biodiversity data repository, subject to completion of data-quality checks and institutional data-sharing procedures. The taxonomic names used in the analyses were standardised against Plants of the World Online and other authoritative taxonomic resources. Additional metadata and analytical scripts will be made available where appropriate to facilitate reproducibility.

## Ethics statement

This study was based primarily on herbarium specimens and published ethnobotanical and medicinal-plant literature. No new interviews with human participants or collection of personally identifiable ethnomedicinal information were undertaken as part of this study. Consequently, the study did not involve direct recruitment or participation of human subjects.

## Author contributions

D.A.Z. conceived the study. D.A.Z., U.A.Y. and I.I. curated and validated the herbarium data, conducted the taxonomic standardisation, performed the analyses, interpreted the results, and prepared the manuscript. All authors contributed to the interpretation of the results, critically revised the manuscript, and approved the final version.

## Preprint statement

This manuscript has been deposited as a preprint and has not undergone peer review. The manuscript is intended for submission to the Journal of Ethnopharmacology.

